# A mediating framework in resting-state connectivity between the medial prefrontal cortex and anterior cingulate in mild cognitive impairment

**DOI:** 10.1101/2024.02.27.582424

**Authors:** Yiyuan Teresa Huang, Sui-Hing Yan, Yi-Fang Chuang, Yao-Chia Shih, Yan-Siang Huang, Yi-Chien Liu, Scott Shyh-Chang Kao, Yen-Ling Chiu, Yang-Teng Fan

## Abstract

Mild cognitive impairment (MCI) is recognized as the prodromal phase of dementia, a condition that can be either maintained or reversed through timely medical interventions to prevent cognitive decline. Considerable studies using functional magnetic resonance imaging (fMRI) have indicated that altered activity in the medial prefrontal cortex (mPFC) serves as an indicator of various cognitive stages of aging. However, the impacts of intrinsic functional connectivity in the mPFC as a mediator on cognitive performance in individuals with and without MCI have not been fully understood. In this study, we recruited 42 MCI patients and 57 healthy controls, assessing their cognitive abilities and functional brain connectivity patterns through neuropsychological evaluations and resting-state fMRI, respectively. The MCI patients exhibited poorer performance on multiple neuropsychological tests compared to the healthy controls. At the neural level, functional connectivity between the mPFC and the anterior cingulate cortex (ACC) was significantly weaker in the MCI group and correlated with multiple neuropsychological test scores. The result of the mediation analysis further demonstrated that functional connectivity between the mPFC and ACC notably mediated the relationship between the MCI and semantic working memory. These findings suggest that altered mPFC-ACC connectivity may have a plausible causal influence on cognitive decline and provide implications for early identifications of neurodegenerative diseases and precise monitoring of disease progression.

## Introduction

Individuals with mild cognitive impairment (MCI) are characterized by having greater cognitive decline than normal aging, while still retaining the ability to perform daily activity (Burns & Zaudig, 2002; Gauthier et al., 2006; Petersen et al., 1999). It has been observed that the prevalence of MCI tends to increase with age (Gillis et al., 2019), and 10-15 % of people with MCI develop dementia each year (Ganguli et al., 2004; Zaudig, 2002). During a prodromal phase of cognitive decline, various brain measurements are employed for early detection. One such example is to monitor amyloid beta proteins (A*β*) in plasma or cerebrospinal fluid over time, serving as a predictor for the maintenance of cognitive health or the development of MCI or Alzheimer’s disease (AD) (Albert et al., 2011; Blasko et al., 2008; Hansson et al., 2010; Jack Jr. et al., 2018; Pesaresi et al., 2006). Furthermore, non-invasive magnetic resonance imaging (MRI) techniques have been widely used, and structural brain volume loss and alterations in blood-oxygen-level dependent signals have been associated with different cognitive stages of aging (Grady, 2000; Pihlajamaki et al., 2009; Ries et al., 2008; Wolf et al., 2004). Notably, resting-state functional MRI is conducted without task requirements or external stimulation, and this feature makes it well-suited for studying brain functional network in patients with different levels of neurological impairment and disability to react to tasks (Park et al., 2011).

Considerable aging MRI research has shown the importance of the medial prefrontal cortex (mPFC) subserving various cognitive processes, such as decision-making (Hosseini et al., 2010; Tisdall & Mata, 2023), memory retrieval (Euston et al., 2012; Grady et al., 2006), and self-referencing of social interaction (Gutchess et al., 2007, 2010). Moreover, the mPFC is a core hub of the default mode network (DMN), which is known for its task-negative functional connectivity and composed of the posterior cingulate cortex, precuneus, and inferior parietal areas (Broyd et al., 2009; Raichle et al., 2001). Previous studies have revealed that the DMN activity decreased as the efficiency of cognitive performance decreased from normal aging, MCI, to AD (Greicius et al., 2003, 2004; Hafkemeijer et al., 2012; Jobson et al., 2021; Raichle, 2015). The disruption of the DMN in aging and neurodegenerative diseases can be explained by the metabolism hypothesis, where a high metabolic rate due to the increased DMN activities subsequently causes more accumulation of Aβ, a protein associated with AD, and thus toxicity within the network (Buckner et al., 2005, 2008). In addition to the large-scale network, mPFC-linked functional connectivity to the subgenual anterior cingulate, the hippocampus, the caudate, and the dorsolateral-prefrontal cortex, has shown weakened in the MCI group, accompanied by inefficient performance in the attention-required and self-inferential tasks (Cai et al., 2017; Liang et al., 2011; Ries et al., 2012; Scherr et al., 2021). Similarly, at the subcortical level, the decreased thalamus-related cortical network, including mPFC and other cortical regions, was found in the MCI group (Z. Wang et al., 2012).

While these correlation findings of mPFC functional connectivity are promising, they do not provide the basis for establishing causal relationships between brain function and cognitive deterioration in patients with MCI. Given that mediation analysis is a valuable tool for investigating the impact of intermediate factors that occur between two variables in a causal relationship, it provides a reliable method for identifying the brain mediators of cognitive changes by utilizing data from functional neuroimaging (Atlas et al., 2010; Cheng et al., 2017; Lindquist et al., 2012). In this study, we conducted a battery of neuropsychological tests and resting-state fMRI scans on both patients with MCI and healthy controls. We hypothesized that poorer cognitive performance and decreased functional connectivity in the mPFC are observed in patients with MCI, compared to healthy controls. Additionally, we anticipated that the variations in resting-state mPFC functional connectivity are linked to differences in cognitive performance. Furthermore, we also performed mediation analyses to examine the causal relationships among MCI, intrinsic brain connectivity, and cognitive alterations.

## Materials and Methods

### Participants

A total of 42 participants with MCI (25 females) and 57 healthy control participants (35 females) were from the Taiwan Precision Medicine Initiative of Cognitive Impairment and dementia (TPMIC) study. In this study, participants assigned to the MCI group were recruited from a memory clinic with a Clinical Dementia Rating (CDR) score of 0.5. They met the 2011 National Institute on Aging–Alzheimer’s Association (NIA-AA) criteria, as confirmed by experienced and certified physicians and a clinical psychologist (Albert et al., 2011). Participants assigned to the healthy control group had no neurological or psychiatric disorders and showed no signs of cognitive decline, as indicated by a CDR score of 0 and a Mini-Mental State Examination (MMSE) score above the cutoff determined by their educational levels (Uhlmann & Larson, 1991). We excluded those with major psychiatric diseases, other neurodegenerative diseases, brain trauma, active cancer, recent hospitalization, and current infection from the two groups. All participants had normal corrected vision and bilateral peripheral hearing during testing. Informed assent and consent were obtained from all participants. All procedures in the present study were approved by the Institutional Review Board of Far Eastern Memorial Hospital (IRB number: FEMH 105147-F) and conducted in accordance with the Declaration of Helsinki.

### General procedures

We performed the CDR test (Morris, 1993) and the MMSE test (Folstein et al., 1975) to measure the global cognitive and functional abilities of each participant. A battery of neuropsychological tests was also used to evaluate various domains of neuropsychological functioning. The tests included forward digit span, symbol substitution, semantic verbal fluency tests of animals, vegetables, fruits, and towns (Isaacs & Kennie, 1973), immediate and delayed logical memory tests (from the third version of the Wechsler Memory Scale) (Wechsler, 1997), the Stroop color-word test (Bondi et al., 2002), as well as part A and B of the color trails tests (CTT) (Reitan & Wolfson, 1985). Note that we replaced English alphabets with colors in part B to exclude biases due to different English levels. We recorded the time duration each participant spent completing each part of the CTT.

### Imaging acquisition

The structural and resting-state fMRI data were acquired on a Skyra 3T MR scanner (Siemens Healthcare, Erlangen, Germany). The scanning procedure included the acquisitions of (1) structural MR images (MPRAGE sequence, field of view (FOV) = 220 × 220 mm, matrix size = 256 × 256, repetition time (TR) = 2060 ms, echo time (TE) = 2.35 ms, flip angle = 10 degrees, and a slice thickness of 0.90 mm with no gap); (2)the resting-state fMRI scans (T2*-weighted GE-EPI sequence, TR = 2000 ms, TE = 24 ms, a total of 180 volumes, 35 axial slices, FOV = 220 × 220 mm, voxel size = 3 × 3 × 4, flip angle = 90 degrees, with a duration 6 minutes and 8 seconds). All participants were instructed to keep their eyes closed and maintain their head position fixed during scanning to obtain high-quality brain images. We collected neuropsychological and imaging data on two separate days (at least one week apart).

### Pre-processing for resting-state fMRI data

Functional resting-state fMRI data was analyzed using CONN toolbox v-21a, a plug-in toolbox in MATLAB-based SPM12 (Wellcome Department of Cognitive Neurology, University College London). The preprocessing pipeline includes functional realignment and unwrap, slice timing correction, outlier detection, co-registration, segmentation, normalization, band-pass filtering, and smoothing (6-mm full width at half maximum Gaussian kernel). Functional images were realigned to correct for head movement, and Artefact Detection Tools were used to scrub outliers. Parameters for realignment and scrubbing were set at an intermediate level (97th percentiles, with linear motion parameters > 0.9 mm and global-signal z > 5 value threshold) for a denoising procedure. All images were segmented into gray matter, white matter, and cerebrospinal fluid. The segmented images were then normalized into standard Montreal Neurologic Institute (MNI) space, with an isotropic voxel size of 1 mm for structural and 2 mm for functional images. Functional images were band-pass filtered from 0.01 to 0.1 Hz.

### Seed-based resting-state functional connectivity analyses

The medial prefrontal cortex (mPFC), defined from the default mode network according to Human Connectome Project (HCP) atlas (Nieto-Castanon, 2020), was chosen as our main region of interest (ROI). The average time series was extracted, and we conducted Pearson’s correlation between target ROI and all other voxels respectively. Fisher’s *z*-score was calculated before being analyzed at the group level. In group-level analysis, we used analysis of variance to examine differences in z scores between normal and MCI participants. Within-group thresholds were set at *p*-uncorrected < 0.001 at the voxel level and *p*-FDR corrected < 0.05 at the cluster level according to Gaussian Random Field Theory (Worsley et al., 1996). Between-group thresholds were set at *p*-uncorrected < 0.001 at the voxel level and *p*-uncorrected < 0.001 at the cluster level.

### Statistical analysis

The chi-square test for categorical variables and independent *t* test for continuous variables were used to compare the demographic characteristics of the two groups. For neuropsychological and resting-state fMRI data, the analysis of variance was used to evaluate the differences between the HC and MCI groups, while age and educational years were set as covariates. To quantify significant differences between the groups, the effect size of Cohen’s *d* and eta squared (*η^2^*) was calculated. Furthermore, we used multiple linear regression analysis to assess the relationship between resting-state functional connectivity and each score of the neuropsychological tests across all participants. Age and years of education were set to be covariates. *P* values (significance set below 0.05) and Adjusted *R^2^* values were evaluated.

### Mediation analysis

If our data show correlations among the following three variables—group, resting-state functional connectivity, and performance on the neuropsychological evaluations, we would conduct the mediation analysis using a Matlab-based toolbox developed by Dr. Wager and colleagues (Wager et al., 2008). Based on the conceptual framework of a mediation effect (Gunzler et al., 2013), we selected the group label (i.e., the HC and MCI groups) as the predictor variable, scores on the neuropsychological tests as the outcome variable, and strengths of mPFC functional connectivity as the mediator variable. Mediation analyses were conducted in a sequential series of steps. The path *a* is to estimate the effects of the predictor variable on the mediator variable; that is, the group difference in strengths of mPFC functional connectivity. The path *b* is to estimate the effect of the mediator variable on the outcome variable while the effects of the predictor variable are controlled. The path *c* is to estimate the effects of the predictor variable on the outcome variable; that is, the total effects of the group on neuropsychological test performance. On the other hand, the path *c’* goes through the mediation variable; that is indirect effect mediated by strengths of mPFC functional connectivity. The *a*b* effect is the mediation effect which is equivalent to *c* – *c’*. In other words, the reduction reflects that path coefficients change, when the predictor-outcome relationship includes the mediator variable. The bootstrap method was used to test statistical significance.

## Results

### Demographic information and neuropsychological evaluations

Table 1 shows demographic and neuropsychological results for the HC and MCI groups. Significant differences were observed in terms of age and educational years. Furthermore, as anticipated, the MCI group exhibited significant inefficiency in all neuropsychological tests, compared to the HC group (all *p* < 0.001), while age and education were controlled (Table 1 and Figure 1). The MCI group had lower scores on the MMSE, logical memory, forward digit span, symbol substitution, verbal fluency, and the Stroop tests, with effect sizes ranging from medium to large. The MCI group also required more time to complete parts A and B of the CTT, with large effect sizes.

**Table 1.** Demographic and neuropsychological results of the study participants.

|  | HC group<br>(N=57) |  | MCI group<br>(N=42) |  | *p* | effect size |
| --- | --- | --- | --- | --- | --- | --- |
|  | Mean | SE | Mean | SE |  |  |
| Age (years) | 65.39 | 0.85 | 74.38 | 1.15 | <.001 | 1.29<sup>a</sup> |
| Education (years) | 11.89 | 0.43 | 8.14 | 0.60 | <.001 | 1.61<sup>a</sup> |
| Sex (F/M) | 35/22 |  | 25/17 |  | .85 | — |
| The MMSE | 28.20 | 0.19 | 20.12 | 0.92 | <.001 | 0.29<sup>b</sup> |
| Logical memory (immediate) | 38.32 | 1.29 | 13.43 | 1.65 | <.001 | 0.47<sup>b</sup> |
| Logical memory (delayed) | 21.86 | 1.01 | 5.25 | 1.22 | <.001 | 0.38<sup>b</sup> |
| Forward digit span | 27.72 | 0.72 | 18.36 | 0.91 | <.001 | 0.27<sup>b</sup> |
| Symbol substitution | 68.45 | 2.24 | 31.65 | 3.01 | <.001 | 0.31<sup>b</sup> |
| Verbal fluency (animals) | 14.33 | 0.39 | 9.16 | 0.76 | <.001 | 0.18<sup>b</sup> |
| Verbal fluency (others) | 38.65 | 0.92 | 21.86 | 1.48 | <.001 | 0.44<sup>b</sup> |
| Stroop test (word) | 94.55 | 2.54 | 67.14 | 3.69 | <.001 | 0.12<sup>b</sup> |
| Stroop test (color) | 80.65 | 2.16 | 52.25 | 2.79 | <.001 | 0.27<sup>b</sup> |
| Stroop test (color-word) | 41.58 | 1.82 | 20.19 | 1.81 | <.001 | 0.27<sup>b</sup> |
| Color trails test A | 50.77 | 4.10 | 118.71 | 10.55 | <.001 | 0.14<sup>b</sup> |
| Color trails test B | 104.04 | 5.28 | 228.45 | 17.07 | <.001 | 0.17<sup>b</sup> |

HC: healthy control; MCI: mild cognitive impairment; SE: standard error; F: female; M: male;
MMSE: Mini-Mental State Examination; superscripted a: the Cohen's *d* effect size; superscripted
b: the eta-squared effect size

**Figure 1.**
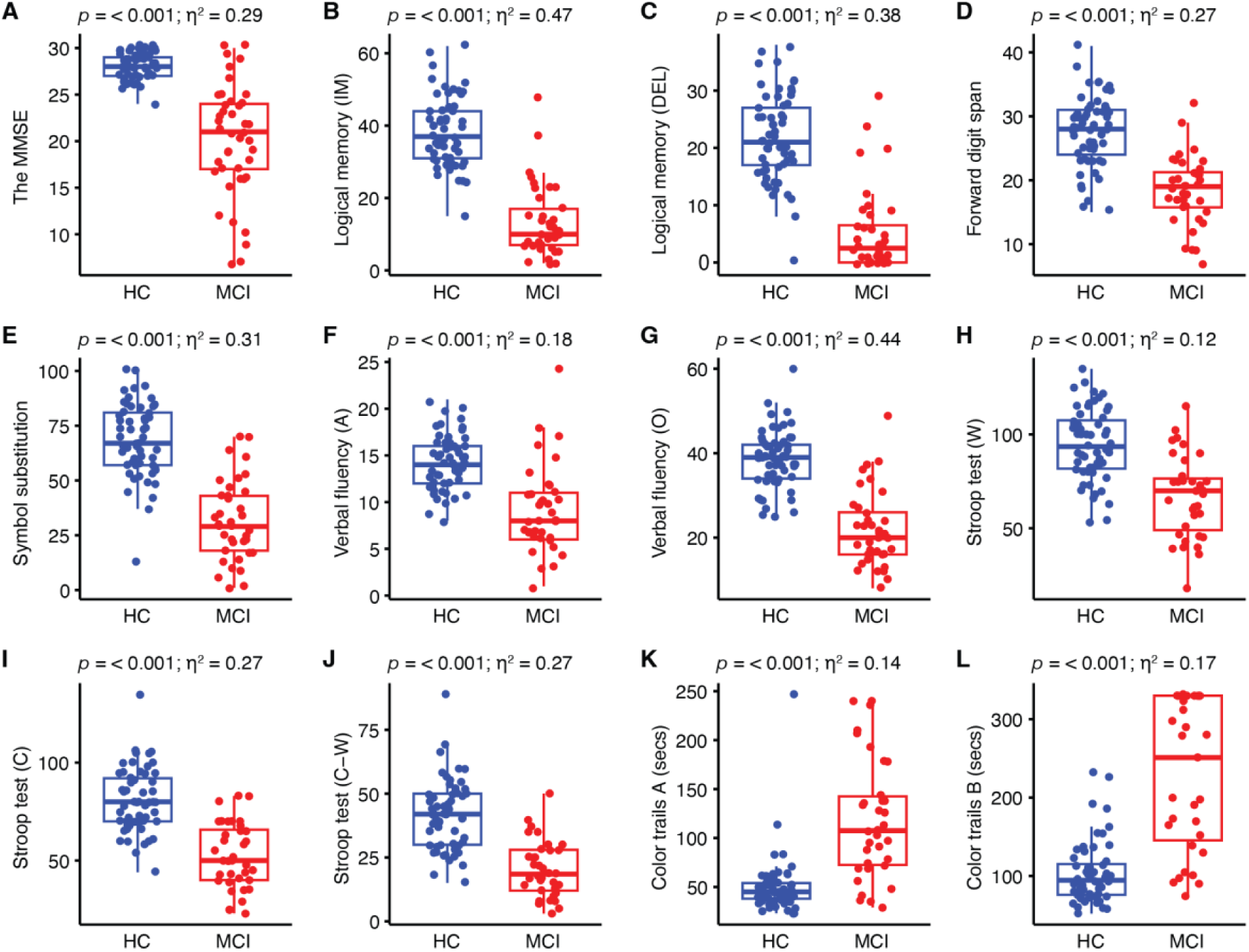
Neuropsychological evaluations in the HC and MCI groups. Differences in neurocognitive tests between the two groups were estimated using ANCOVA analysis and the method of eta squared (*η^2^*) with age and educational years controlled. Immediate and delayed subtests in the logical memory test are denoted as IM in panel B and DEL in panel C, respectively. The verbal fluency subtests for animal and other categories are denoted as A in panel F and O in panel G, respectively. The Stroop subtests for word, color, and both are denoted as W in panel H, C in panel I, and C-W in panel J, respectively. Dots represent individual values. The unit in panels K and L is second.

### Resting-state fMRI results

#### Within-group functional connectivity

In the HC group, we found significant positive functional connectivity between the mPFC and the fusiform gyrus, parahippocampal gyrus, cerebellum, and cortical areas, including the superior frontal gyrus, posterior cingulate, angular gyrus, inferior temporal gyrus, middle frontal gyrus, and superior temporal gyrus (Figure 2A and Supplementary Table S1). In the MCI group, the mPFC exhibited positive connectivity to the fusiform gyrus, lentiform nucleus, cerebellum, and cortical areas including the superior frontal gyrus, posterior cingulate, angular gyrus, middle temporal gyrus, and inferior frontal gyrus (Supplementary Table S2 and Figure 2B). Additionally, negative functional connectivity was observed between the mPFC and the caudate, fusiform gyrus, brainstem, cerebellum, and cortical areas including the inferior temporal gyrus, superior parietal lobule, inferior parietal lobule, middle frontal gyrus, precentral gyrus, middle temporal gyrus, superior frontal gyrus, and lingual gyrus in HC group (Figure 2A and Supplementary Table S1). In the MCI group, the mPFC exhibited negative connectivity to the caudate, cerebellum, and cortical areas including the inferior parietal lobule, superior parietal lobule, precentral gyrus, middle frontal gyrus, superior frontal gyrus, and inferior frontal gyrus (Figure 2B and Supplementary Table S2).

**Figure 2.**
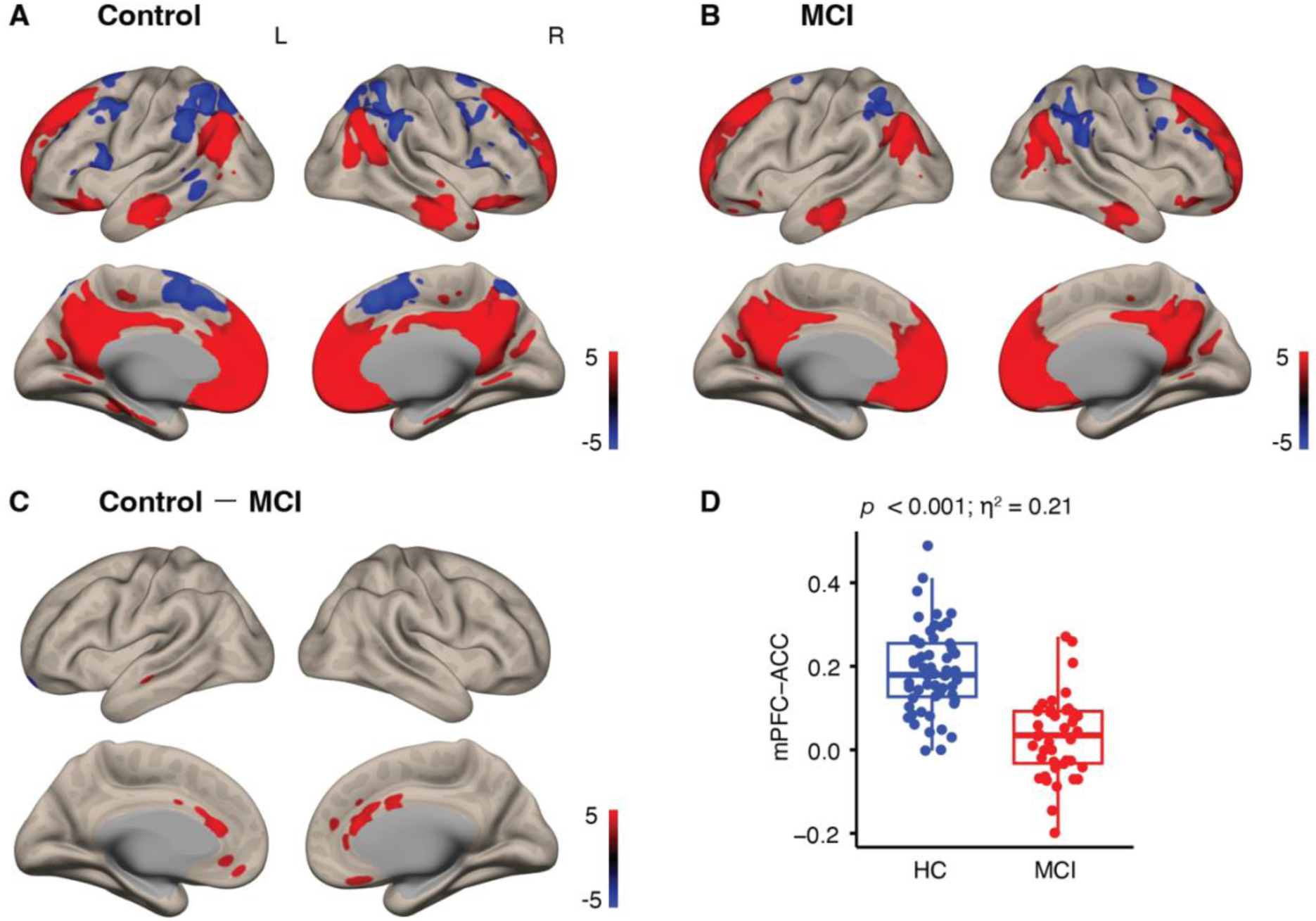
mPFC-specified functional connectivity in the HC and MCI groups. For the within-group analyses, significant functional connectivity across participants in the HC and MCI groups is respectively shown in panel A and panel B. For the between-group analyses, significant differences between functional connectivity in the HC and MCI groups (HC – MCI) are shown in panel C. Individual strengths of mPFC-ACC functional connectivity were extracted and compared in terms of the group, shown in panel D.

#### Between-groups functional connectivity

Table 2 and Figure 2C show significant differences where stronger mPFC functional connectivity was found in the HC group than in the MCI group. Specifically, there is functional connectivity between the mPFC and the areas including the anterior cingulate cortex (ACC), left medial frontal gyrus (MFG), left middle temporal gyrus (MTG), right caudate, right mid-anterior cingulate (mACC), and right mPFC. In Figure 2D, individual strengths of functional connectivity between the mPFC and ACC were extracted, and the significant group difference is shown (*p* < 0.001, *η^2^* = 0.21).

**Table 2.** Regions showing significant differences in resting-state functional connectivity between the HC and the MCI groups.

| Brain area | MNI Coordinates |  |  | T value | Cluster size<br>(voxels) |
| --- | --- | --- | --- | --- | --- |
|  | x | y | z |  |  |
| ***HC group > MCI group*** |  |  |  |  |  |
| Anterior cingulate | 0 | 32 | 18 | 5.92 | 328 |
| Medial frontal gyrus | -4 | 40 | -12 | 4.61 | 53 |
| Middle temporal gyrus | -50 | -18 | -18 | 4.46 | 31 |
| Caudate | 2 | 10 | 8 | 4.36 | 35 |
| Mid-anterior cingulate | 4 | 6 | 30 | 4.23 | 30 |
| Medial prefrontal gyrus | 12 | 46 | 18 | 3.87 | 56 |
| ***MCI group > HC group*** |  |  |  |  |  |
| None |  |  |  |  |  |

### Correlations between brain functional connectivity and neuropsychological performance

Then, we focused on functional connectivity showing significant differences between the groups (i.e., mPFC-ACC, mPFC-MFG, mPFC-MTG, mPFC-caudate, mPFC-mACC, and mPFC-mPFC). The linear regression analysis was performed to estimate associations between each functional connectivity and each neuropsychological test. Here, in Figure 3, we show strengths of mPFC-ACC functional connectivity increase as higher scores (i.e., better performance) on the MMSE, immediate and delayed logical memory scores, forward digit span scores, symbol substitution scores, verbal fluency scores, and the Stroop tests scores. Also, the strengths of mPFC-ACC functional connectivity increase as shorter completion time for parts A and B of the color trial tests. Results of associations between each of the rest significant functional connectivity and each neuropsychological test are shown in Figure S1.

**Figure 3.**
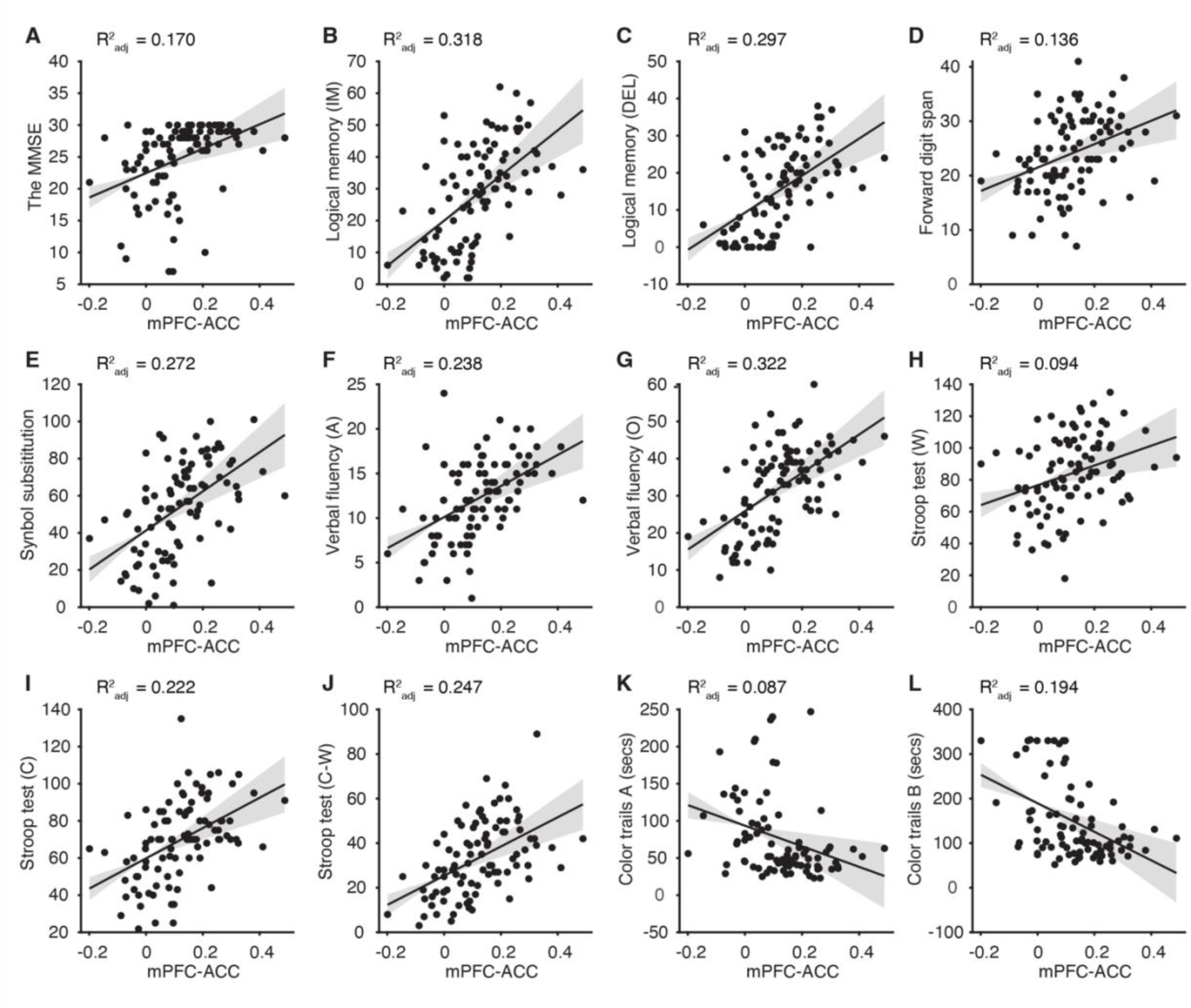
Regressions of functional connectivity against neuropsychological evaluations. Strengths of mPFC-ACC functional connectivity were assigned as an independent variable, and the scores of 12 cognitive tests were assigned as a dependent variable respectively for each linear regression model. Dots represent individual values. Solid lines represent predictions based on the strengths of mPFC-ACC functional connectivity. Gray shades represent the range within 97.5% confidence intervals.

### Mediation analysis results

We observed (1) the significant group difference in the neuropsychological tests, (2) the significant group difference in functional connectivity, and (3) the significantly positive association between stronger functional connectivity and better performance on the tests. Given these significant statistical results, we then performed the mediation analysis with bootstrap resampling. We found that mPFC-ACC functional connectivity plays a significant mediator in the relationship between the group and the animal verbal fluency performance (Figure 4). This suggested that decreased functional connectivity between the mPFC and ACC in MCI may lead to a greater decline in performance on the animal verbal fluency test (*a* = -0.16, SE = 0.02, *p* < 0.001; *b* = 8.50, SE = 3.51, *p* = 0.02; *c* = -5.02, SE = 0.84, *p* < 0.001; *c’* = -3.62, SE = 1.01, *p* < 0.001; *a*b* = -1.39, *Z* = -2.63, *p* = 0.01). Non-significant results were found from the analyses with each of other significant functional connectivity as the mediator and each of the other neuropsychological tests as the outcome.

**Figure 4.**
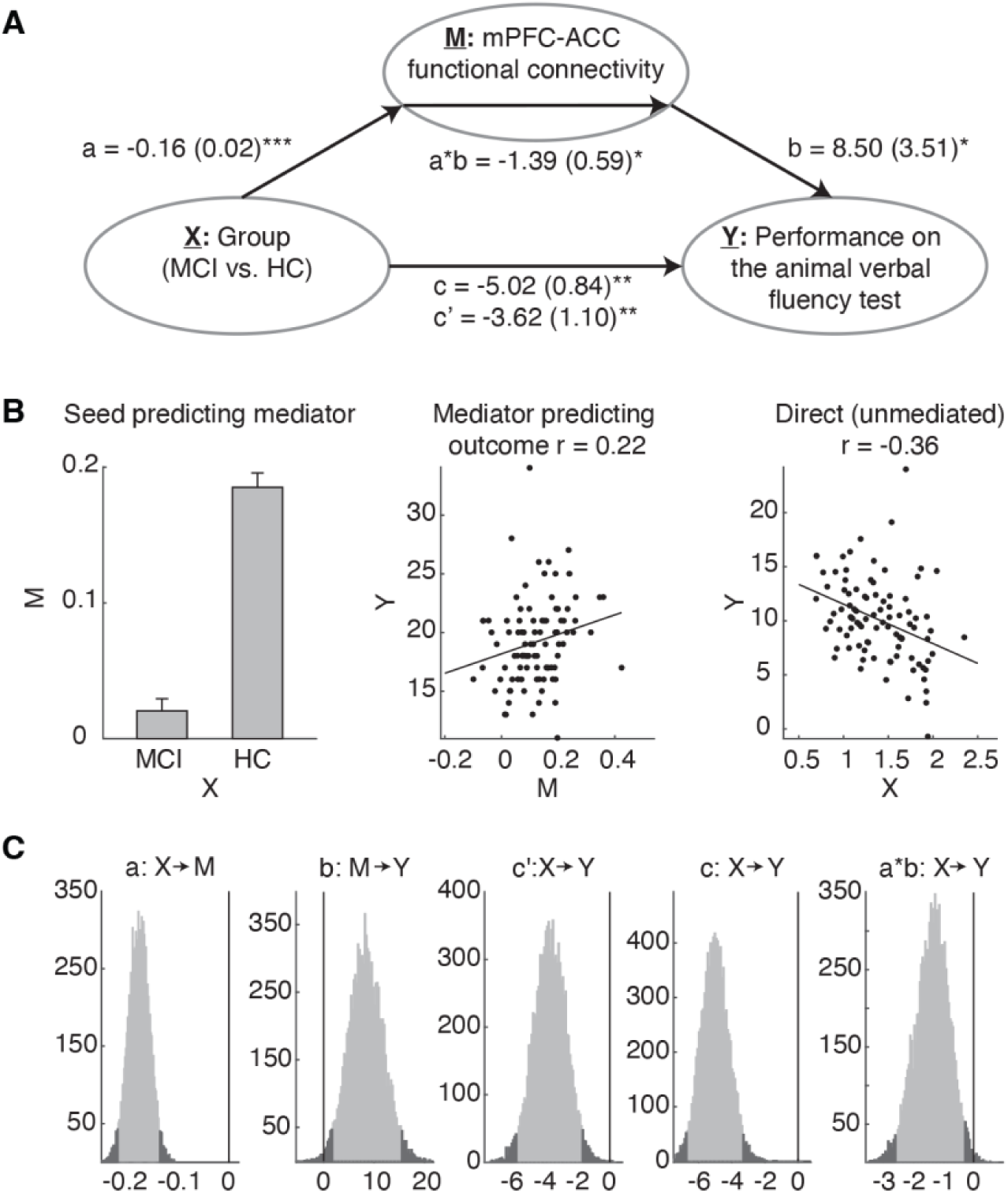
Mediation analysis results. The path diagram in panel A shows the group label as the predictor variable (denoted as X), strengths of mPFC-ACC functional connectivity as the mediator variable (denoted as M), and scores on the animal verbal fluency test as the outcome variable (denoted as Y). The paths *a* and *b* represent the relationships between X and M and between M and X, respectively. The path *a*b* represents the mediation effect. The paths *c* and *c’* respectively represent the total and indirect effects of the group on animal verbal fluency performance, and the latter was evaluated when the mediator variable was considered. The path coefficients and their standard errors were labeled above the paths. The relationships in the paths *a*, *b*, and *c* are visualized in panel B. The bootstrap method was used for each path to define 97.5% confidence intervals in histograms shown in panel C (two-tailed). \*\**p* < 0.001; \**p* < 0.05.

## Discussion

The current study provides evidence for linkages among the MCI group, intrinsic brain connectivity, and impaired cognitive functions. Behaviorally, patients with MCI had significantly poorer performance on multiple cognitive measures, compared to the healthy control group. Neurophysiological findings indicated that there was a notable attenuation of functional connectivity between the mPFC and ACC in the MCI group and correlations between functional connectivity strengths and neuropsychological scores. Furthermore, the results of the mediation analysis showed that the attenuated mPFC-ACC functional connectivity causally influences cognitive performance in semantic working memory in the MCI group.

Using the mPFC as a seed of interest, we suggested that functional connectivity to the ACC serves as a mediator on the pathology of MCI. The ACC, situated in the frontal part of the cingulate cortex, is widely recognized for its role in cognitive and motor processes, such as error detection, attention allocation, and motor preparation (Cameron S. Carter et al., 1999; Luks et al., 2002; Turken & Swick, 1999). The causal impact of atypical brain activations on cognitive impairment, suggested by our mediation analysis results, was further supported by lesion studies. The compelling evidence has shown that people with damages in the ACC and mPFC were unable to modulate their behavioral responses when conflict with previous stimulus presentations appeared (di Pellegrino et al., 2007; Modirrousta & Fellows, 2008; Swick & Jovanovic, 2002). However, it stands in contrast with our mediation analysis results where the Stroop test, one of the conflict tasks, was not significantly mediated by weak mPFC-ACC connectivity in the MCI. We interpret the incongruency with two possible reasons: (1) relatively minor deficits in conflict processes in MCI and (2) compensations from other brain areas. It has been found that MCI patients might have the partially intact ability to respond to congruent (i.e., non-conflict) versus incongruent (conflict) trials, where a lower accuracy with a similar reaction time or a similar accuracy with a longer reaction was observed, compared to healthy controls (Bélanger et al., 2010; Borsa et al., 2018; P. Wang et al., 2013). Furthermore, increased activities of brain areas including the ACC, prefrontal, and inferior parietal cortices were observed in response to the conflict task in the MCI group than in the HC group (Li et al., 2009), suggesting the preservation in MCI through a compensatory network.

Despite the lack of valid causality between decreased mPFC-ACC connectivity and the conflict task, the neural influence on verbal fluency showed significance. It has been widely reported that verbal fluency is a linguist marker to detect people at an early stage of cognitive decline or with subjective cognitive impairment which could hardly be verified by standard tests (Ostberg et al., 2005). Although language-specialized brain areas notably include the inferior frontal gyrus (IFG), and superior and middle temporal gyrus (Friederici, 2011), the domain-general brain areas such as cingular-opercular network are also involved in language processes (Fedorenko & Thompson-Schill, 2014; Geranmayeh et al., 2014; Saur et al., 2006). One study showed that the dorsal ACC (dACC) and pre-supplementary motor area were correlated with recovery of language functions after four months of left-hemisphere brain stroke (Geranmayeh et al., 2017). Furthermore, given the semantic verbal fluency tasks that require producing names of one category (e.g., animal) as many as possible, retrieval of one’s knowledge and experience is also one of the key skills. Compared to speech delivery, individuals with MCI may be more prone to memory retrieval, and thus, the mPFC is preferentially activated. To be more cautious, we also admit that mPFC-ACC functional connectivity could not fully account for the cause of poor semantic verbal fluency in MCI. Other possibilities include a hub area not appropriately allocating domain-general brain areas (e.g., dACC) and language-specialized brain areas (e.g., IFG), or disconnection between the IFG and dACC.

Although our findings provide a better understanding of mPFC-ACC connectivity as a causal mediator in cognitive impairment, there are still some advances to be achieved in future research, including addressing methodological issues, analyzing directed functional connectomes, using structural equation modeling (SEM) as a stricter statistical approach, and incorporating non-invasive brain stimulation. First, the matching of MCI and HC groups exhibited significant differences (i.e., age and educational years).

Although we incorporated these factors as covariates in the data analyses, we cannot disregard the possible influence they may have on the conclusion. Second, these findings were obtained in a cross-sectional study, indicating the necessity for a longitudinal investigation to elucidate potential links between mPFC-ACC functional connectivity and disease progression. Thirdly, in the current study, we correlated the activity of the mPFC as a seed with the whole brain area without the assumption of directional influence from one area to the other area. Indeed, this non-directed functional connectivity is easily included in the clinical protocol for fast screening or help of diagnosis, but it fails to extract the etiology of dysfunctional connectomes. A combination of analysis of directed connectomes in future research enables further dissection; for example, determining whether weak mPFC-ACC connectivity results from the inadequately activated mPFC connecting to the ACC or vice versa.

Moreover, the SEM is different from the standard regression approach we used to capture the casual relationship, and it allows for the inclusion of multiple independent variables, mediators, or outcomes (Gunzler et al., 2013). For example, two hypothesized connections (e.g., one language-specific and the other domain-general) both act as mediators. Furthermore, compared to estimating regression models separately step by step, causal relationships in the SEM are verified by simultaneously considering direct and indirect effects, based on a conceptual model and postulated paths. However, the SEM is sensitive to the number of model parameters (Tomarken & Waller, 2003, 2005). Thus, to hypothesize the plausible number of mediators and/or outcomes, the standard regression approach could work as a safeguard procedure. Also, better verifications of a concept model and hypothesized pathways require enough data samples which should be considered in future research. Furthermore, with statistical approaches (i.e., mediation analysis), we can quantify possible causal connectomes on cognitive impairment. To further merge the results into the interventional protocol, without doubt, the causality should be supported by experimental manipulation. There are two common stimulations, transcranial direct current stimulation (tDCS) and transcranial magnetic stimulation, used to activate or deactivate an activity of a target brain area. It has been reported that the tDCS applied over the prefrontal cortex and combined with physical therapies can improve verbal fluency in MCI patients with Parkinson’s disease (Manenti et al., 2016); however, the relevant findings remain inconsistent (Birba et al., 2017; Elder & Taylor, 2014). We hope that focal stimulations on the presumed target brain network with reference to the statistical-causal results could reconcile the inconsistency and improve the effectiveness and efficiency of treatment in MCI.

## Conclusion

This study demonstrated alterations in cognitive functioning and patterns of brain functional connectivity in MCI patients. Importantly, our investigation constructed a mediating framework and highlighted the causal link between altered mPFC-ACC connectivity and semantic working memory deficits among MCI patients. The significance of these findings not only contributes valuable insights into the neurobiological underpinnings of aging-related cognitive changes but also has practical implications for early detection, disease monitoring, and the effective treatment protocol in clinical trials for neurodegenerative conditions.

## Supporting information

All Supplementary Tables and Figures

## Author contributions

S-H.Y., Y-F.C., Y-L.C., and Y-T.F. conceived and conceptualized the study. Y.T.H., S-H.Y., Y-F.C, Y-C.S., S.S-C.K., Y-S.H., Y-C.L., and Y-T.F. contributed to acquisition, analysis, or interpretation of data for the work. Y.T.H., and Y-T.F. conducted the necessary literature reviews and drafted the first manuscript. S-H.Y., Y-F.C., Y-C.S., Y-S.H., Y-C.L., and Y-L.C. provided critical feedback and helped shape the manuscript. All authors approval of the final version to be submitted.

