## Supplementary material for "A mediating framework in resting-state connectivity between the medial prefrontal cortex and anterior cingulate in mild cognitive impairment": All Supplementary Tables and Figures

### SUPPLEMENTARY MATERIALS

**Table S1**. Regions showing significantly positive and negative functional connectivity in the HC group

| Brain area | MNI Coordinates | | | *T* value | Cluster size (voxels) |
|---|---|---|---|---|---|
| | x | y | z | | |
| ***Positive correlation*** | | | | | |
| Superior frontal gyrus | −4 | 58 | −2 | 27.37 | 15081 |
| Posterior cingulate | −4 | −52 | 24 | 17.13 | 6001 |
| Angular gyrus | −44 | −66 | 36 | 12.1 | 1747 |
| Inferior temporal gyrus | −62 | −12 | −18 | 9.59 | 969 |
| Fusiform gyrus | 60 | −6 | −22 | 9.4 | 976 |
| Angular gyrus | 52 | −62 | 34 | 9.0 | 1192 |
| Middle frontal gyrus | 34 | 34 | −14 | 7.1 | 466 |
| Cerebellar tonsil | −6 | −54 | −46 | 6.88 | 330 |
| Parahippocampal gyrus | −24 | −20 | −18 | 6.61 | 389 |
| Parahippocampal gyrus | 26 | −20 | −20 | 6.39 | 170 |
| Middle frontal gyrus | −28 | 32 | −16 | 6.03 | 387 |
| Superior temporal gyrus | 42 | 24 | −30 | 5.08 | 133 |
| Cerebellar tonsil | 46 | −60 | −44 | 4.99 | 139 |
| Inferior semi-lunar lobule | −46 | −66 | −42 | 4.09 | 43 |
| ***Negative correlation*** | | | | | |
| Inferior temporal gyrus | 40 | −12 | −22 | 6.76 | 42 |
| Medial frontal gyrus | 8 | 24 | 48 | 6.52 | 1309 |
| Superior parietal lobule | −38 | −54 | 56 | 6.48 | 1661 |
| Inferior parietal lobule | 36 | −46 | 42 | 6.25 | 1031 |
| Middle frontal gyrus | −46 | 2 | 48 | 5.90 | 311 |
| Uvula | −34 | −66 | −26 | 5.85 | 83 |
| Middle Frontal Gyrus | 44 | 42 | 28 | 5.64 | 199 |
| Precentral gyrus | −44 | 8 | 10 | 5.34 | 255 |
| Pyramids | 24 | −66 | −28 | 5.33 | 79 |
| Pyramids | −18 | −66 | −30 | 5.32 | 50 |
| Inferior semi-lunar lobule | 20 | −78 | −46 | 5.27 | 108 |
| Middle frontal gyrus | 46 | 6 | 56 | 5.18 | 76 |
| Middle temporal gyrus | −54 | −48 | 2 | 5.14 | 160 |
| Superior frontal gyrus | −42 | 36 | 32 | 4.92 | 45 |
| Middle frontal gyrus | 44 | 16 | 18 | 4.71 | 173 |
| Superior parietal lobule | 16 | −66 | 56 | 4.62 | 235 |
| Caudate | −16 | −32 | 18 | 4.52 | 64 |
| Lingual gyrus | −30 | −64 | 0 | 4.34 | 43 |
| Declive | 8 | −76 | −22 | 4.10 | 52 |

**Table S2**. Regions showing significantly positive and negative functional connectivity in the MCI group

| Brain area | MNI Coordinates | | | *T* value | Cluster size (voxels) |
|---|---|---|---|---|---|
| | x | y | z | | |
| ***Positive correlation*** | | | | | |
| Superior frontal gyrus | −6 | 56 | −2 | 19.61 | 12935 |
| Posterior cingulate | 6 | −52 | 28 | 11.94 | 4304 |
| Angular gyrus | −46 | −66 | 34 | 7.59 | 1213 |
| Middle temporal gyrus | 60 | −2 | −24 | 6.75 | 465 |
| Angular gyrus | 50 | −66 | 36 | 6.65 | 879 |
| Inferior frontal gyrus | −32 | 30 | −12 | 6.57 | 167 |
| Fusiform gyrus | −58 | −18 | −22 | 6.51 | 417 |
| Inferior frontal gyrus | 32 | 28 | −14 | 6.08 | 179 |
| Inferior semi-lunar lobule | −48 | −68 | −42 | 5.11 | 120 |
| Inferior semi-lunar lobule | 46 | −64 | −44 | 4.68 | 69 |
| Lentiform nucleus | −12 | 2 | 6 | 4.25 | 56 |
| ***Negative correlation*** | | | | | |
| Inferior parietal lobule | 36 | −46 | 44 | 7.26 | 949 |
| Caudate | 2 | 12 | 10 | 5.95 | 91 |
| Superior parietal lobule | −34 | −52 | 54 | 5.39 | 470 |
| Precentral gyrus | 34 | 4 | 36 | 5.08 | 47 |
| Middle frontal gyrus | 44 | 46 | 24 | 4.99 | 200 |
| Middle frontal gyrus | 22 | 2 | 60 | 4.95 | 100 |
| Superior frontal gyrus | −26 | 0 | 72 | 4.63 | 52 |
| Superior parietal lobule | 12 | −68 | 58 | 4.37 | 56 |
| Uvula | 26 | −72 | −24 | 4.33 | 44 |
| Inferior frontal gyrus | 58 | 14 | 30 | 4.12 | 75 |

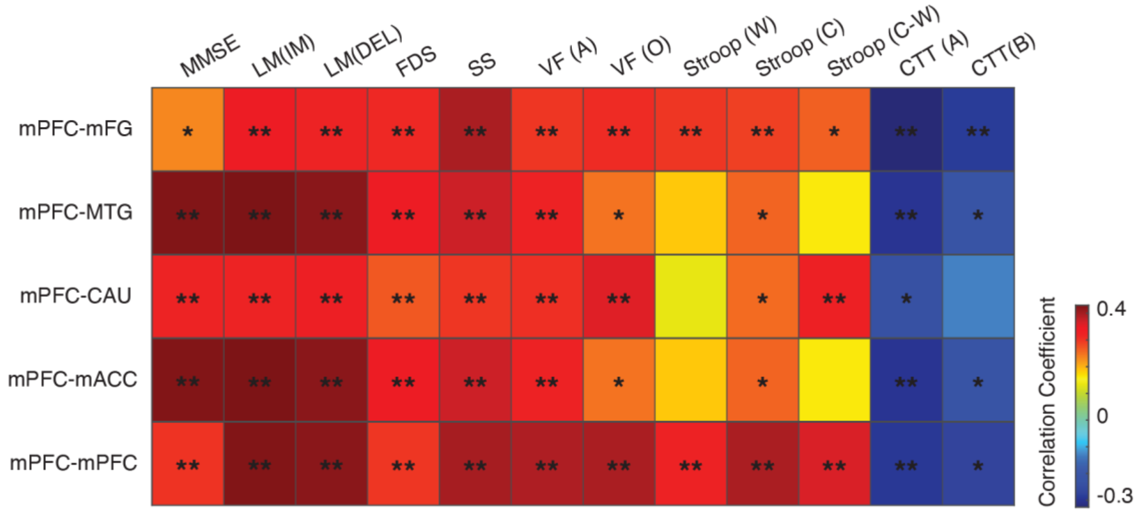


**Figure S1**. **Heatmap showing correlation coefficients between brain functional connectivity and neuropsychological evaluations**. Warm and cold colormaps respectively represent positive and negative correlations between each connectivity (listed in the top row) and each evaluation (listed in the top column). $^{**}p < 0.01$, $^{*}p < 0.05$.

Abbreviations: mFG, medial frontal gyrus; MTG, middle temporal gyrus; CAU, caudate; mACC, mid-anterior cingulate; LM (IM), immediate logical memory tests; LM (DEL), logical memory (delayed recall); FDS, forward digit span; SS, symbol substitution; VF (A), verbal fluency tests of animals; VF (O), verbal fluency tests of others categories (vegetables, fruits, and towns); Stroop (W), the Stroop word test; Stroop (C) the Stroop color test; Stroop (C-W), the Stroop color and word test; CTT (A), part A of the color trails test; CTT (B), part B of the color trails test.
